# Limb-Selective Regions in the Lateral Temporal Lobe Shrink from Childhood to Adulthood

**DOI:** 10.64898/2026.03.06.709762

**Authors:** Selina Cohnen, Larissa Kahler, Seong Dae Yun, Kerstin Konrad, Marisa Nordt

**Author notes:** Corresponding author: Selina Cohnen, Pauwelsstraße 19, 52074 Aachen.

## Abstract

Perceiving hand gestures and inferring others’ actions and emotions from movements of hands and limbs play an important role in every-day interactions, especially in young children. The perception of categories such as limbs, bodies, or faces is supported by category-selective regions in the temporal lobe. Some category-selective regions, such as those reacting selectively to body parts or limbs, exist both on the ventral and on the lateral side of the temporal lobe, and are part of the ventral and lateral stream, respectively. While it was recently shown that limb-selective regions in the ventral stream shrink from childhood to adulthood, the developmental trajectory of limb-selective regions in the lateral stream remains unknown. To close this gap in knowledge, we acquired functional magnetic resonance imaging (fMRI) data in 21 children aged 10 – 12 years and 20 adults while they watched images of 10 visual categories including limbs and whole bodies. We first replicate the decrease of limb-selectivity from childhood to adulthood in the ventral temporal lobe. Across several analyses, our results further demonstrate that limb-selective regions in the lateral temporal lobe shrink as well, particularly in the left hemisphere. Underlining the specificity of our finding, we show that lateral body-selective regions show no significant development from childhood to adulthood. These findings advance our understanding of the developmental trajectories of limb- and body-selective regions and of the ventral and lateral visual streams more broadly.

## 1. Introduction

Long before children can speak fluently, they rely on their hands to guide others’ attention to objects of interest and to interpret the actions of those around them by watching their hands. For instance, around 9–12 months, infants understand and follow pointing gestures to interpret others’ communicative intentions and goals, and soon begin pointing themselves to create joint attention (Carpenter et al., 1998; Liszkowski et al., 2004; Tomasello et al., 2007). Likewise, in adulthood, hands and the gestures hands produce convey a wealth of socially relevant information about other individuals. This includes the actions they perform, the emotional states they are in or their intentions and thoughts. These examples highlight the importance of the perception of body parts, such as hands and limbs, for everyday social interactions.

High-level visual regions in the temporal lobe are critical for the perception of body parts (Downing et al., 2001; Peelen & Downing, 2005). In fact, the temporal lobe contains functional regions responding preferentially to visual categories including body-parts or limbs (Downing et al., 2001; Peelen & Downing, 2005; Weiner & Grill-Spector, 2011), faces (Andrews & Ewbank, 2004; Kanwisher et al., 1997; Puce et al., 1998), places (Epstein & Kanwisher, 1998) and words (Dehaene et al., 2002). These regions are referred to as *category-selective regions,* because they show a stronger response to a specific category compared to other types of visual input. Critically, regions selective for some of these categories are present both in the ventral and the lateral temporal lobe and are thought to be part of the ventral and lateral visual stream, respectively. The ventral stream extends from primary visual cortex (V1) to ventral temporal cortex (VTC) and supports perception and recognition (Grill-Spector et al., 2000; Moutoussis & Zeki, 2002; Tong et al., 1998), whereas the lateral stream extends from V1 via areas involved in motion processing to the superior temporal sulcus (STS) and is thought to be involved in dynamic and social processing (Allison et al., 2000; Gandolfo et al., 2024; Gomez et al., 2019; Hein & Knight, 2008; Pitcher & Ungerleider, 2021; Weiner & Grill-Spector, 2013) and action processing (Wurm & Caramazza, 2022). The body-(part) selective region in the ventral stream is the fusiform body area (FBA; Peelen & Downing, 2005), located in the occipito-temporal sulcus (OTS). In the lateral stream, the body-(part) selective region has been termed the extrastriate body area (EBA; Downing et al., 2001) consisting of three body-selective patches surrounding the motion selective region hMT+ (Weiner & Grill-Spector, 2011).

How do these regions develop? Research on the development of body-selective regions has yielded mixed findings: One cross-sectional study reported increases in body-selectivity from childhood to adulthood in both ventral and lateral body-selective regions (Ross et al., 2014). In contrast, another cross-sectional study found no significant development of the FBA (Peelen et al., 2009), although the size of nearby face-selective regions increased across the same age range (Aylward et al., 2005; Cantlon et al., 2011; Golarai et al., 2007, 2010; Nordt et al., 2021; Peelen et al., 2009; Scherf et al., 2007). Results of this latter study were replicated by a recent longitudinal study showing that ventral regions responding selectively to whole bodies and limbs follow different developmental trajectories throughout childhood and adolescence (Nordt et al., 2021). Specifically, body-selective regions showed no developmental change, whereas limb-selective regions decreased in size from 5 to 17 years. Interestingly, limb selectivity in VTC decreased with age, while selectivity for words increased in the same region. These prolonged developments throughout childhood and adolescence suggest that neural resources originally dedicated to limb processing are repurposed to word recognition, a developmental mechanism described as “cortical recycling” (Dehaene & Cohen, 2007; Nordt et al., 2021).

These results raise the question how limb-selective regions in the lateral stream develop from childhood to adulthood. Prior research suggests two hypotheses: One is that lateral limb-selective regions decrease in size with age, similar to the previously observed decrease in the ventral stream (Nordt et al., 2021). While it is unknown what is driving this development, a possible explanation might be that changes in limb-selectivity are linked to changing viewing behaviour for limbs (Kubota et al., 2024): That is, the visual diet – the distribution of visual inputs an individual sees in daily life – regarding the frequency of hands may change throughout childhood. In fact, studies using head-mounted cameras to record what children view, showed that hands occur highly frequently in the visual experience of infants and toddlers (Frank et al., 2012; Jayaraman et al., 2017; Yoshida & Smith, 2008). In addition, recent eye-tracking studies indicate that viewing behaviour towards hands develops from childhood to adulthood: Preschool children allocate more visual attention to limbs when viewing naturalistic scenes, whereas literate adults preferentially attend to text (Linka et al., 2023, 2025). Such developmental changes in viewing behaviour may influence limb-selective responses in the ventral and lateral temporal lobe, leading to similar developmental trajectories across the ventral and lateral streams.

An alternative hypothesis suggested by prior literature is that lateral limb-selective regions remain stable. This view is supported by findings from Golarai and colleagues (2007), who showed that ventral face-selective regions expand from childhood to adulthood, whereas no such effects were found for lateral regions. In addition, the ventral and lateral stream differ with regard to the processing of static and dynamic stimuli (Pitcher et al., 2019), and these differences may affect the development of the two streams. Taken together, these results indicate that category-selective regions in the ventral and lateral stream follow different developmental trajectories and suggest that lateral limb-selective regions may show no developmental changes.

The decrease of limb-selectivity in the ventral stream also raises the question whether tool-selective regions (Bracci et al., 2012; Chao et al., 1999) may show a similar development. Notably, much like limb-selective regions, tool-selective regions have been reported both in the ventral and lateral temporal lobe: The ventral tool-selective region is located in the medial part of the fusiform gyrus (Bracci et al., 2016), the lateral tool-selective region is located in the left lateral occipitotemporal cortex (Peelen et al., 2013; Pillet et al., 2024). Two aspects suggest that tool-selective regions may undergo a developmental trajectory similar to that of limb-selective regions: First, tools, much like hands, manipulate objects and may often be viewed at the same time. Second, lateral tool-selective regions partially overlap with nearby located hand-selective regions in the brain (Bracci et al., 2012; Pillet et al., 2024). In fact, while a prior developmental study reported no significant development of tool-selectivity from childhood to adulthood (Dekker et al., 2011), they showed that children had additional ventral tool-selective clusters in the left fusiform gyrus compared to adults.

To address these questions, we collected fMRI data in 10-12-year-old children and adults and measured the size of their limb-selective regions in the ventral and lateral streams.

## 2. Methods

### 2.1 Statement on ethical regulations

This study was approved by the Ethics Committee of the RWTH Aachen.

### 2.2 Participants

Healthy participants with normal or corrected-to-normal vision and hearing were recruited from local institutions. Exclusion criteria were the diagnosis of a psychiatric or neurological disorder and/or a family history of dyslexia, and contraindications for MRI measurements, such as metallic implants, as assessed by a short non-standardized questionnaire.

21 children aged 10 – 12 years (11 female; mean age: *M* = 11.3 years; SD = 0.83) and 20 adults aged 22 – 61 years (17 female; mean age: *M* = 35.1 years; SD = 11.6) participated in this study. The age range for children was chosen because previous research demonstrated significant development in high-level visual regions from age 10 to adulthood (Golarai et al., 2010; Meissner et al., 2019; Nordt et al., 2019, 2021). The initial child sample consisted of 23 children (12 female; *M* = 11.31 years; SD = 0.84), but data of two children were excluded (one due to technical problems, one due to excessive head motion). All children reported German as their native language and two children spoke additional languages. 19 adults reported German as their native language, and one participant reported Russian as their native language but was fluent in German. Four children and two adults were left-handed as assessed by the Edinburgh Handedness Inventory (Oldfield, 1971).

While adults participated in one session only, children took part in two, which were conducted within a one-month period. This second session was introduced to compensate for possible data loss due to motion. If children already completed two runs with high scan quality in the first session (see below, **Quality control of MRI Data)**, the runs of the first session were included in the analysis. Overall, each session lasted around 90–120 minutes. During the first session, children participated in a mock scanner training (see below, **Mock-Scanner Training**). Otherwise, the MRI scanning procedure was the same in both sessions. To prevent fatigue, behavioural testing was distributed over the two timepoints for the children. Adults participated in additional measurements as part of a larger study. Parents gave written consent and children gave verbal and/or written assent, adults gave written consent. Children were rewarded with a 10 Euro bookstore voucher and small toys of their choice for each session. Adults received 60 Euro

### 2.3 Behavioural testing

Outside the scanner participants performed a standardized reading test (Salzburger Lese-und Rechtschreibtest II, SLRT-II; Moll & Landerl, 2010) in which they were instructed to read as many real words (task 1) and pseudowords (task 2) as possible in 60 seconds. All participants had scores in the normal range.

### 2.4 Mock-Scanner Training

Children were prepared for their first MRI session with an extensive mock-scanner training to get familiarized with the scanner environment in a child-friendly way and to reduce motion during scanning (Raschle et al., 2012). During the training, children were introduced to scanner noises and practiced lying still. The importance of remaining still was underlined using examples of blurred versus sharp images (Raschle et al., 2012). The duration of the mock-scanner training was adapted to individuals’ needs and typically lasted between 20 and 30 minutes.

### 2.5 MRI acquisition parameters

Structural and functional MRI data were collected at the Forschungszentrum Jülich on a 3-Tesla Siemens scanner using a 64-channel head coil. A high-resolution T1-weighted structural MRI scan was acquired for anatomical imaging. Structural imaging producing T1 contrast was acquired over a period of approximately 10 minutes with the following parameters: 2300ms TR, 2.32ms TE, 8° flip angle, 0.9mm³ voxel size (isotropic). Additionally, a short T1-weighted anatomical “inplane scan” with identical slice orientation to the functional data was collected to facilitate alignment between the anatomical and the functional images with the software package mrVista (see below). Functional imaging (see below) comprised three separate runs, each lasting 5 minutes and 24 seconds. Imaging parameters of the functional sequences are 1000ms TR, 28.00ms TE, 60° flip angle, 2.5mm³ voxel size leading to a slice thickness of 2.5mm.

### 2.5 fMRI paradigm: functional localizer

Participants performed three runs of a modified version of the fLoc-localizer (Stigliani et al., 2015). This modified version contains stimuli of five visual domains. Each domain includes two categories: faces (child faces, adult faces), bodies (headless bodies, limbs), places (corridors, houses), words (real words, pseudowords), and tools (shovels, pens; **Fig. 1A**). Specifically, we made the following changes to the stimuli of the original localizer: We used real German words instead of numbers and a different set of pseudowords, matched to the German real words. Further, we used tools instead of objects, and our limb stimuli did not contain any feet or legs, but only hands (some also showing arms). The remaining stimuli are the same as reported in prior publications (Gomez et al., 2019; Nordt et al., 2019, 2021, 2023; Stigliani et al., 2015). As in the original version of the localizer, stimuli were grey-scaled images and presented in 4s blocks at a rate of 2 Hz. During scanning, participants were instructed to fixate on a small fixation cross and to perform an odd-ball task during which they were supposed to press a button whenever a scrambled image appeared.

**Figure 1.**
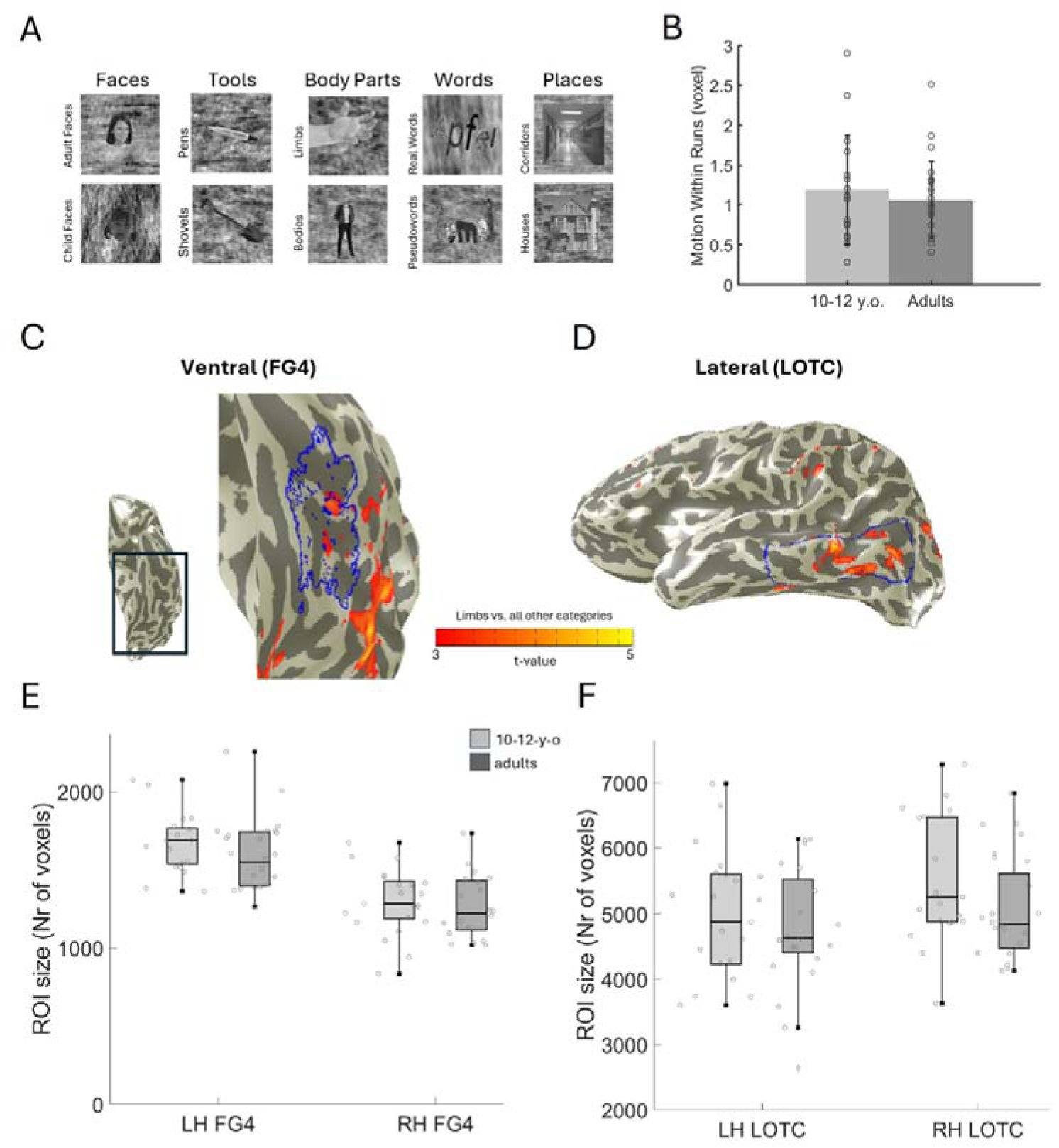
Experimental Stimuli, Quality Control and Example ROIs. **A** Stimuli presented to the participants during MRI scanning comprise five domains (adapted from Stigliani et al., 2015): faces, tools, body parts, words, and places. Each domain includes two categories: child faces and adult faces, pens and shovels, headless bodies and limbs, real words and pseudowords, and houses and corridors. The faces illustrated were not presented in the real experiment and are the faces of two of the authors (M.N. and S.C), one of them as a child. **B** Mean within-run motion values (in voxels) during the included functional MRI runs for children (in light grey) and adults (in dark grey). **C** The ventral cytoarchitectonic ROI FG4 is outlined in blue on the ventral temporal cortex in the left hemisphere of a 10-year-old participant. The ventral limb-selective activation (defined by a t-value > 3 on the voxel-level for the contrast limbs vs. all other categories) falls within the FG4 ROI. **D** The anatomically defined LOTC ROI is outlined in blue on the lateral side of the left temporal lobe of an 11-year-old participant. The lateral limb-selective activation (defined by a t-value > 3 on the voxel-level for the contrast limbs vs. all other categories) falls within the LOTC ROI. **E** Sizes of the anatomical FG4 ROIs for children and adults in voxels. The horizontal line in each box denotes the median value. Whiskers extend to the most extreme data points that do not qualify as outliers. **F** Same as E, but for LOTC ROIs.

### 2.6 Quality control of MRI Data

Data were excluded if participants moved their head more than three functional voxels within a run or between two runs, as in prior publications (Dalski et al., 2024). As not all participants retained all three runs after applying this criterion, only two runs per participant were included in the analysis to ensure consistency across the samples. If all three runs were valid, the first two runs were chosen, as we expected participants to be more alert during the two initial runs. Applying these criteria, the average motion of included runs did not differ significantly between the groups (**Fig. 1B**).

### 2.7 Processing of MRI data

For structural data, we used FreeSurfer’s (https://surfer.nmr.mgh.harvard.edu) “recon-all” pipeline to generate a cortical surface reconstruction of each participant. For the functional data, analysis was performed in MATLAB R2019a (MathWorks, Inc.) using the mrVista software package (https://github.com/vistalab/vistasoft). Functional MRI data was aligned to each participant’s structural data in its’ native space. Motion correction was performed within and between functional runs. No spatial smoothing and no slice-timing correction were applied. Prior work using similar pipelines has adopted this strategy to preserve high spatial resolution and to avoid blurring of responses to different categories across anatomical boundaries (Gomez et al., 2017; Nordt et al., 2021, 2023). We transformed the time courses into percentage signal change by dividing each voxel’s data by the average response across the entire run. To estimate the contribution of each of the 10 conditions, a general linear model (GLM) was fit to each voxel by convolving the stimulus presentation design with the hemodynamic response function (HRF). We used the HRF as implemented in SPM (https://www.fil.ion.ucl.ac.uk/spm/).

### 2.8 Anatomical Regions of interest (ROIs)

#### Cytoarchitectonic regions of interest

To examine the development of limb-selectivity in the ventral stream, we leveraged the finding that the ventral limb-selective region consistently falls into the cytoarchitectonic region FG4 (Weiner et al., 2017). Cytoarchitectonic regions are characterized based on how their cells are organized. While they are identified in post-mortem tissue, a probabilistic estimate of the cytoarchitectonic areas of the ventral temporal lobe can be obtained from an atlas (Rosenke et al., 2018). This atlas includes the cytoarchitectonic regions FG1, FG2, FG3 and FG4 in VTC (Caspers et al., 2013; Lorenz et al., 2015). For this study, we focused on the FG4 ROI, since the ventral limb-selective region falls into this part of the brain (**Fig. 1C**). This relationship has been validated on children’s brains previously (Kubota et al., 2023).

#### Lateral occipitotemporal cortex (LOTC)

To examine the development of limb-selectivity in the lateral stream, we defined lateral occipitotemporal cortex (LOTC) ROIs based on anatomical landmarks, as in prior studies (Bugatus et al., 2017; Weiner & Grill-Spector, 2011) on the inflated cortical surface of each hemisphere in each participant (**Fig. 1D**). This ROI was chosen because prior research showed that the lateral limb-selective region falls into this anatomical ROI (Weiner & Grill-Spector, 2013). The inferior border of the LOTC ROI was defined by the lateral border of the OTS, the superior border by the inferior side of the superior temporal sulcus (STS). The anterior border of the LOTC ROI was defined by the anterior tip of the mid fusiform sulcus (MFS), the posterior boundary was set posterior to the lateral occipital sulcus (LOS).

### 2.9 Definition of category-selectivity and category-selective regions

To examine the development of category-selectivity, we applied two approaches: First, an observer-independent approach to examine the number of voxels selective to a given category within anatomically defined ROIs (FG4 and LOTC ROIs, see above, **Cytoarchitectonic regions of interest; Lateral occipitotemporal cortex**) in each participant. Second, we manually defined functional ROIs in each participant and examined their sizes. We used the same definition for category-selectivity at the voxel level across approaches: We contrasted the responses for a given category to those for all other categories (for instance, limbs vs. all other categories). A voxel was classified as category-selective for a threshold of t>3 for the given contrast. This threshold was chosen because previous studies showed that category-selective regions can be defined reliably in children and adults using this threshold (Gomez et al., 2017; Natu et al., 2016; Nordt et al., 2021). In addition, to validate that our results do not depend on this specific threshold, we also conducted a threshold-independent analysis (see below).

#### Definition of category-selectivity in the anatomical ROIs

We used a data-driven and observer-independent approach to examine how category-selectivity develops in high-level visual cortex as in prior publications (Nordt et al., 2021). We assessed the selectivity to each category in anatomically defined ROIs (ventral FG4 and lateral LOTC ROIs) in each participant and counted the number of voxels that passed this threshold. We report both the relative number of category-selective voxels relative to the overall number of voxels within anatomical ROIs as well as absolute numbers of selective voxels (as supplemental analyses). Additionally, we conducted a threshold-independent analysis. For this, we calculated the t-value for a given contrast for each voxel in an anatomical ROI and then examined the mean t-value across all voxels.

#### Functional regions of interest (fROIs)

We further manually delineated lateral limb-selective regions in both hemispheres on each participant’s native cortical surface by using a combination of functional and structural information as in previous studies (Nordt et al., 2021; Weiner & Grill-Spector, 2011). Again, we used a threshold of t>3 (voxel-level) for the definition of functional ROIs and the contrast “limbs vs. all other categories”. The lateral limb-selective region typically includes three limb-selective patches forming a crescent-shaped activation pattern around the motion-selective area hMT+ (Weiner & Grill-Spector, 2011). Specifically, it consists of one activation patch on the lateral occipital sulcus/middle occipital gyrus located posterior to hMT+, the second patch on the middle temporal gyrus located anterior to hMT+ and the third patch on the inferotemporal gyrus located inferior to hMT+. This limb-selective region overlaps partially with the extrastriate body area (EBA; Downing et al., 2001), which is typically defined using whole body stimuli.

### 2.10 Statistical analyses

Statistical analyses were performed using MATLAB R2019a (MathWorks, Inc.). To determine whether there were significant differences between the groups, we first performed Levene’s test to control for the assumption of homogeneity of variances. If homogeneity of variances was present, we performed independent-samples t-tests, if not, Welch’s t-test was performed. Levene tests and results for all categories are reported in the supplements. Normal distribution was assumed, since the sample sizes for each group were N=20 or larger. Significant effects were controlled for multiple comparisons (across the 10 categories in each ROI) using the Benjamini-Hochberg-FDR correction as implemented in MATLAB R2019a (MathWorks, Inc.).

## 3. Results

### No group differences in the amount of motion during scanning or in the sizes of anatomically defined ROIs

We first tested whether children and adults differed significantly regarding their head motion during scanning. Independent samples t-tests revealed no significant differences between the two groups (t(39)=0.712, p=0.48) (**Fig. 1B**). Thus, differences in functional activation are unlikely to stem from motion differences. Next, we performed independent-samples t-tests to determine whether children and adults differed significantly in the sizes of the anatomically defined ROIs. There was no significant difference across groups for the FG4 ROIs (lh: t(39)=1.06, p =0.29; rh: t(39)=0.52, p=0.61, **Fig. 1E**) and no difference for the LOTC ROIs (lh: t(39)=0.73, p=0.47; rh: t(39)=0.83, p=0.41, **Fig. 1F**).

### Replication of the decrease of limb-selectivity in the ventral stream

We first aimed to replicate previous findings reporting a decrease in limb-selectivity in the ventral temporal lobe (Nordt et al., 2021). Given that the ventral limb-selective region falls into the FG4 ROI (Kubota et al., 2023; Weiner et al., 2017), we tested whether the proportion of voxels within this ROI responding preferentially to limbs relative to other stimuli, differed between children and adults. Our results revealed that children had more limb-selective voxels compared to adults in the FG4 ROI in the left (t(39)=2.63, p=0.01, FDR-corrected p=0.05, Welch’s t-test, **Fig. 2A**) and the right hemisphere (t(39)=2.09, p=0.04), although the latter effect did not survive correction for multiple comparisons for the 10 categories (FDR-corrected p=0.21). In contrast, for whole bodies, no statistically significant differences between the groups in either the left (t(39)=0.75, p=0.46, Welch’s t-tests) or the right (t(39)=0.67, p=0.51) hemisphere were found, in line with prior results (Nordt et al., 2021; Peelen et al., 2009).

**Figure 2.**
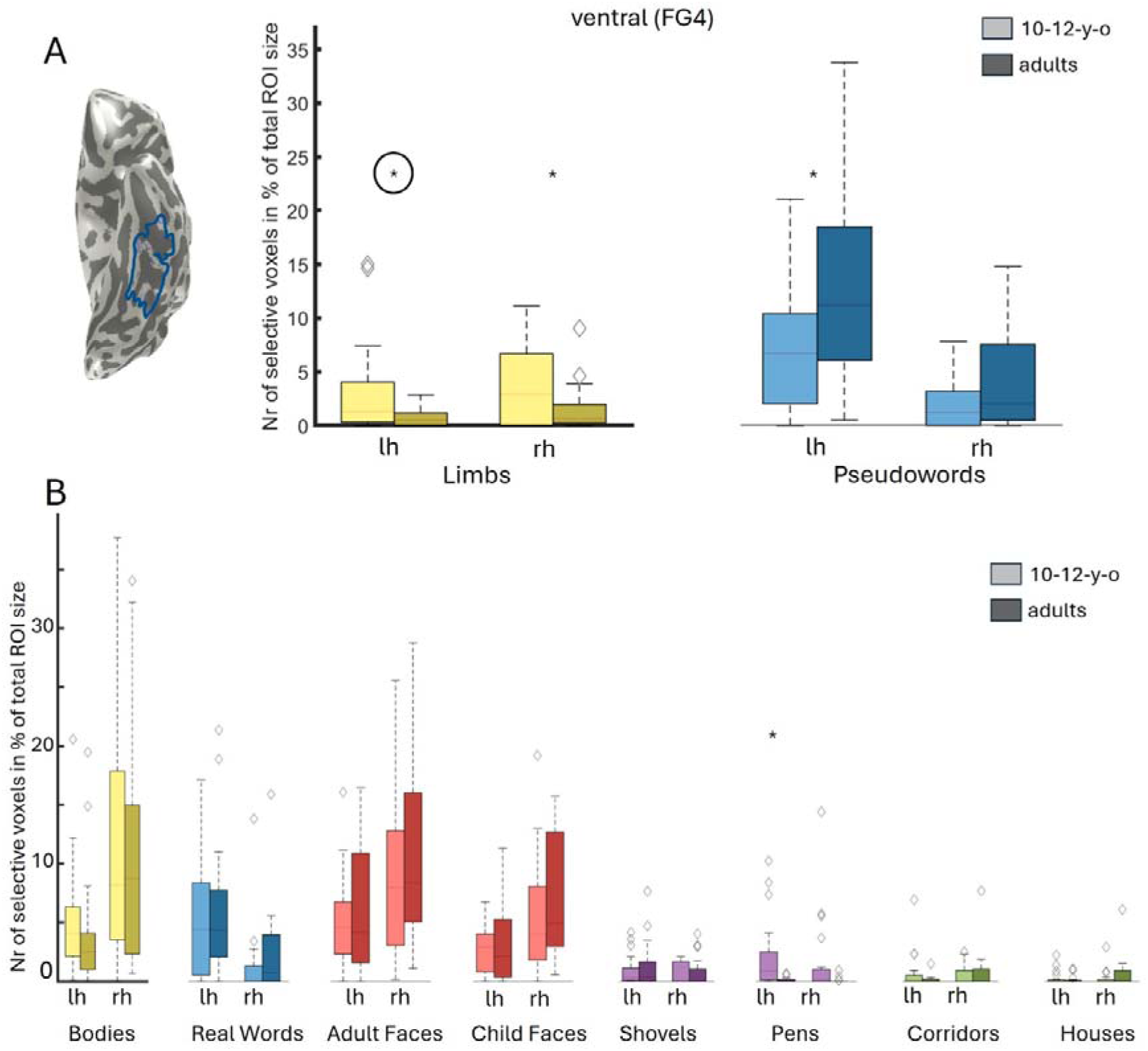
Development in ventral ROIs (FG4) **A** Left: Example showing the FG4 ROI in a 10-year-old participant. Right: Relative number of limb- and pseudoword-selective voxels in the ventral cytoarchitectonic FG4 ROI in 10-12-year-old children (brighter colours) and adults (darker colours). The horizontal line in each box denotes the median value. Whiskers extend to the most extreme data points that do not qualify as outliers. Diamonds are outliers, defined as lying beyond 1.5 interquartile ranges from the first or third quartile. Black asterisks indicate significant differences across the age groups, circles around asterisks indicate significant effects after correction for multiple comparisons for the 10 categories. **B** Same as A but for the categories bodies, real words, adult faces, child faces, shovels, pens, corridors and houses. Categories belonging to the same domain are displayed in the same colour.

Since previous research revealed that decreases in limb-selectivity in VTC are linked to increases in pseudoword-selectivity (Nordt et al., 2021), we next examined the development of voxels selective to real- and pseudowords. Mirroring previous findings, we found a significant increase of the proportion of pseudoword-selective voxels in the FG4 ROI from childhood to adulthood in the left hemisphere (t(39)=-2.54, p=0.02, FDR-corrected p=0.07) which was trending after FDR-correction, but not in the right hemisphere (t(39)=-1.83, p=0.08, **Fig. 2A**). Surprisingly, for real words, we found no significant development in neither the left (t(39)=-0.185, p=0.86) nor the right (t(39)=-0.938 p=0.35) hemisphere (**Fig. 2B**).

Interestingly, selectivity to pens, one of the tool categories, showed a similar trajectory to limb-selectivity. In fact, our results revealed a significant decrease of the number of voxels selective to pens from childhood to adulthood in the left (t(39)=2.87, Welch’s t-test p=0.01, FDR-corrected p=0.05) but not the right FG4 ROI (t(39)=1.98, Welch’s t-test p=0.06, FDR-corrected p =0.22; **Fig. 2B**). No such development was observed for shovels, the other tool category (lh: t (39)=-0.67, p=0.51; rh: t(39)=-0.33, p=0.74). Because ventral tool-selective regions lie in the medial fusiform gyrus (Bracci et al., 2016), these regions should be mainly located in the FG3 ROI. We therefore performed the same analysis in FG3 and found a significant decrease for pens (t(39)=2.6, p=0.017, Welch’s t-test) and for shovels (t(39)=2.2, p=0.035, Welch’s t-test) in the left but not in the right hemisphere (pens: t(39)=1.58, p=0.13, Welch’s t-test; shovels: t(39)=1.63, p=0.12, Welch’s t-test) **(Fig. S1).** Thus, it is possible that the decrease in the number of pen-selective voxels in the left FG4 ROI is driven by tool-selective regions in FG3 that partially overlap with the FG4 ROI. Notably, the overall number of tool-selective voxels in both ROIs was very small.

We found no significant effect for any of the other categories (**Fig. 2B; Tables S1, S2, S3)**, including faces. This is in line with previous research because the FG4 ROI captures mFus-faces (FFA-2), but not pFus-faces (FFA-1), and only pFus-faces is expected to develop within the tested age range (Nordt et al., 2021).

### Selectivity to limbs decreases in the lateral stream

Next, we examined whether and how limb-selective regions develop in the lateral stream by using two methodological approaches: First, we manually delineated lateral limb-selective functional ROIs in each hemisphere of each participant. Second, we applied the same observer-independent approach as in the ventral stream: Here, we counted the proportion of limb-selective voxels relative to the overall number of voxels in anatomically defined LOTC ROIs. While the first approach captures the development of clustered limb-selective activation, the second approach (i) may also include non-clustered limb-selective activation, (ii) is observer-independent and (iii) serves as an internal replication.

We first examined the development of manually delineated lateral limb-selective regions which could be defined in 19 children in the left and in 20 in the right hemisphere, and in 20 adults in the left and 18 in the right hemisphere.

**Figure 3A** displays the lateral view of the left hemispheres of the ten children (upper row) and ten adults (lower row) with the highest number of limb-selective voxels in the left lateral temporal lobe. Qualitative inspection of these maps suggests that children exhibit larger lateral limb-selective regions than adults (**Fig. 3A**; images of all participants can be found in **Fig. S2**). We next turned to quantitative analyses: This revealed larger limb-selective regions in children compared to adults both in the left (t(37)=3.753, p=0.0004, significant at the Bonferroni-corrected threshold (p=0.0008) and right hemispheres (t(38)=2.54, p=0.01, Bonferroni-corrected p=0.0245; **Fig. 3B**). The difference in the region’s size between groups was large: In children the left lateral limb-selective region had a mean size of 718 voxels (Median=598, SD=350), in adults this size was reduced to half the size with 357 voxels (Median=329, SD=212).

**Fig. 3.**
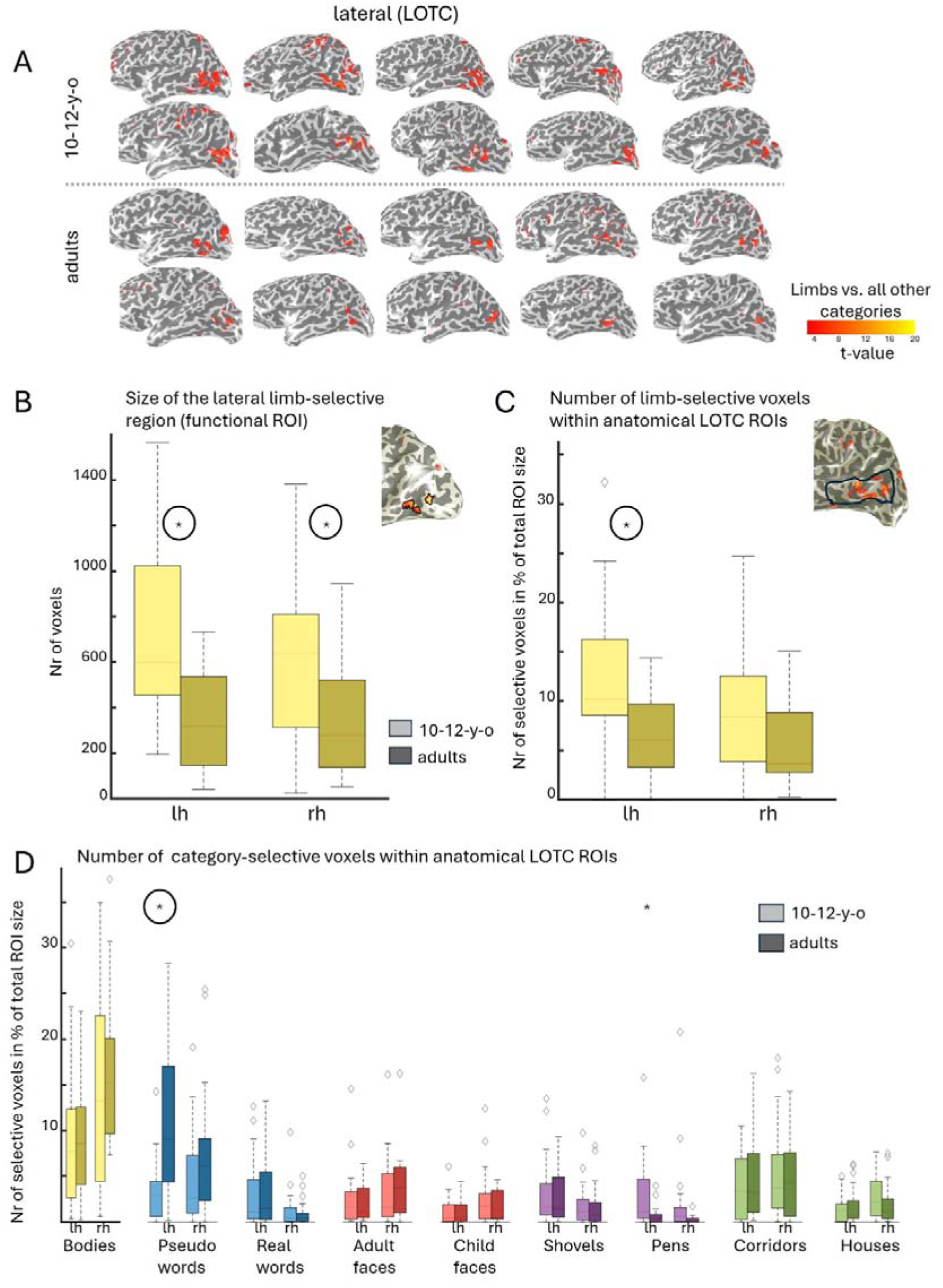
Development in lateral ROIs (LOTC) **A** The 10 children (upper rows) and adults (lower rows) with the highest number of limb-selective voxels in LOTC. Red patches depict limb-selective activation (limbs vs. all other stimuli, t>3). Images of all participants can be found in the supplements. **B** Boxplots: Size of functional limb-selective ROIs (children: N=19 in lh; N=20 in rh; adults: N=20 in lh; N=18 in rh) in the lateral temporal lobe, measured in the number of voxels; Inset: Example for a lateral limb-selective region in the left hemisphere of an 11-year-old participant. **C** Boxplots: Relative number of limb-selective voxels out of all voxels in the left and the right anatomically defined LOTC ROI (children: N=21 in lh and rh; adults: N=20 in lh and rh); Inset: Example showing the limb-selective activation falling into the left LOTC ROI of an 11-year-old participant. **D** Same as boxplots in C but for bodies, pseudowords, real words, adult faces, child faces, shovels, pens, corridors and houses. Across all panels: Values of children are displayed in brighter colours and adults in darker colours. The horizontal line in each box denotes the median value. Whiskers extend to the most extreme data points that do not qualify as outliers. Diamonds are outliers, defined as lying beyond 1.5 interquartile ranges from the first or third quartile. Black asterisks indicate significant differences across the age groups, circles around asterisks indicate significant effects after correction for multiple comparisons for the 10 categories.

In the second methodological approach, we counted the proportion of limb-selective voxels within the LOTC ROIs. Results revealed a significant difference in the number of limb-selective voxels in the left hemisphere (t(39)=2.74, p=0.01, FDR-corrected p=0.04; **Fig. 3C**), with children exhibiting a larger number of limb-selective voxels compared to adults. In the right hemisphere, this effect was only trending and did not survive correction for multiple comparisons (t(39)=2.00, p=0.05, FDR-corrected p=0.39; **Fig. 3C**; all values in **Tables S4, S5, S6**). We repeated these analyses using the absolute number of limb-selective voxels within LOTC ROIs for internal replication, revealing a significant decrease in the left (t(39)=2.88, p=0.01, FDR-corrected p=0.056) and right hemispheres (t(39)=2.29, p=0.03, FDR-corrected p=0.139 for rh; **Tables S7, S8, S9)** that do not survive FDR correction.

To confirm that our results do not depend on a certain t-value used to determine category-selectivity, we also performed a threshold-independent analysis and compared the mean limb-selectivity across all voxels in the LOTC ROIs between the two groups. This analysis yielded a significant decrease of mean limb-selectivity from childhood to adulthood in the left (t(39)=2.97 p=0.01, FDR-corrected p=0.05), but not the right LOTC ROI (t(39)=1.42, p=0.16; **Fig. S3; Tables S10, S11, S12**). In sum, these analyses show that limb-selective regions in the lateral stream shrink from childhood to adulthood and that this effect is more pronounced in the left compared to the right hemisphere.

As prior results demonstrated that the decrease of limb-selectivity in the ventral stream is directly linked to increases in selectivity to pseudowords (Nordt et al., 2021), we next tested whether the decrease in the number of limb-selective voxels in the lateral stream is paralleled by an increase in the number of voxels selective to another category. In particular, we reasoned that word-selective regions in the lateral temporal lobe, located on the inferior occipital sulcus (Stigliani et al., 2015; Zhan et al., 2023), may expand with development and partially overlap with our LOTC ROIs. In fact, we found that adults had a significantly larger number of voxels selective to pseudowords compared to children in the left (t(39)=-3.96, p=0.0003, FDR-corrected p=0.006, Welch’s t-test, **Fig. 3D**) but not right LOTC (t(39)=-1.42, p=0.16, Welch’s t-test, **Fig. 3D**). This effect was specific to pseudowords, as no significant differences between the groups were observed for real words (lh: t(39)=-0.45, p=0.65; rh: t(39)=0.55, p=0.58, **Fig. 3C**).

The parallel decrease in limb-selectivity and increase in pseudoword-selectivity in left LOTC raises the question whether these two developments may be linked as proposed for the ventral stream, where limb-selective regions are thought to be repurposed into word-selective regions as children learn to read (Nordt et al., 2021). While this link can only be examined in longitudinal data, we approached this question indirectly by testing whether the development of selectivity to pseudowords and limbs in LOTC were linked to reading ability. To do so, we first tested whether performance on a reading test (assessing the number of words read in one minute) is linked to the number of voxels selective to pseudowords in left LOTC. As expected, this analysis yielded a significant positive correlation between reading score and the number of pseudoword-selective voxels in the left hemisphere (r=0.45, p=0.003), even after controlling for age (r_p_=0.349, p=0.03; **Fig. S4A**). Next, we reasoned that if the decrease in selectivity to limbs in the lateral stream was linked to reading ability, there should be a negative correlation between the number of limb-selective voxels in LOTC and reading ability that is irrespective of age. While we found a significant negative correlation between the number of voxels selective to limbs and the number of words read within one minute (r=-0.449, p=0.003; **Fig. S4B**), this correlation was only trending after controlling for age (age-controlled r_p_=-0.287, p =0.07).

### Specificity of the decrease of limb-selectivity in the lateral stream

Finally, we tested the specificity of the decrease of limb-selectivity in the lateral stream. We asked whether the decrease in selectivity extends to two closely related categories including (i) whole body stimuli and (ii) tools, which are manipulated by hands. To do so, we first compared the relative number of body-selective voxels between the groups. Our results show no significant differences across age groups in either the left (t(39)=-0.34, p=0.19) or the right (t(39)=-1.05, p=0.96) hemisphere (**Fig. 3D**), suggesting that body- and limb-selective regions in the lateral temporal lobe follow different developmental trajectories. We next tested the development of the number of tool-selective voxels in the lateral stream. While we found a significantly larger number of voxels selective to pens in children compared to adults in the left hemisphere, this effect did not survive correction for multiple comparisons (t(39)=2.6, Welch’s t-test p=0.01, FDR-corrected p=0.09; **Fig. 3D**) and was not significant in the right hemisphere (t(39)=1.82, Welch’s t-test p=0.08, FDR-corrected p=0.31). No significant development was observed for the number of selective voxels for shovels from childhood to adulthood (**Fig. 3D**). We observed no significant development for any of the other categories **(Tables S5, S6).**

## 4. Discussion

Our study has three main findings: First, we replicated the previously reported decrease of limb-selectivity in the ventral stream from childhood to adulthood (Nordt et al., 2021). Second, we showed that this decrease also occurs in the lateral stream, where it was more pronounced in the left compared to the right hemisphere. Critically, this result was observed across different methodological approaches: (i) a functional ROI approach, (ii) an observer-independent approach, where we counted the number of limb-selective voxels within anatomically defined ROIs, and (iii) a threshold-independent approach, where we examined the mean t-value for limbs across all voxels in these anatomically defined ROIs. While the decrease of limb-selectivity in the left hemisphere was significant across all approaches, the decrease in the right hemisphere was only significant in the functional ROI approach. Third, we showed that across the ventral and lateral streams, the development of selectivity to limbs differed from that to whole bodies, corroborating prior findings (Nordt et al., 2021; Peelen et al., 2009).

In the following paragraphs we discuss possible reasons for the decrease of limb-selectivity, the link to the development of representations to other categories, potential behavioural ramifications, the specificity of this finding and implications of this result for the development of visual streams, as well as the timing of the observed developmental effects.

While it is currently unknown what drives the decrease of limb-selectivity across the ventral and lateral streams, our findings are in line with previous behavioural studies using eye-tracking to investigate differences in gaze behaviour between children and adults (Linka et al., 2023, 2025). These studies demonstrated that children look more at hands when freely viewing natural complex scenes, compared to adults, who look more at text. Thus, a possible explanation linking our findings with these behavioural results is that changes in viewing behaviour towards limbs from childhood to adulthood may affect category-selective responses to these stimuli in both the ventral (Kubota et al., 2024) and lateral stream, leading to a decrease of limb-selectivity. This hypothesis also suggests that adults who frequently use and look at their hands (e.g. surgeons or artists), might show increased limb-selectivity compared to adults in other professions.

A related question raised by our results is whether the observed changes in limb-selectivity might be linked to other developmental changes in the brain. Longitudinal results have revealed that decreases in limb-selectivity in the ventral stream were directly linked to increases in word-selectivity (Nordt et al., 2021) in line with the idea of a recycling of category-selectivity (Dehaene et al., 2010; Dehaene & Cohen, 2007; Nordt et al., 2021). In this context, the decrease of limb-selectivity, and the parallel increase of pseudoword-selectivity observed in the lateral temporal lobe, raise the question whether these developments are linked as well. Consistent with this idea is the lateralization of the observed effects. Our results demonstrate a stronger and more consistent decrease of limb-selectivity in the left compared to the right LOTC, matching the left-lateralized increase in selectivity to pseudowords. While a limitation of our study is that we are missing longitudinal data to determine whether cortical recycling also occurs in the lateral stream, we approached this question indirectly by correlating reading ability with the number of limb-selective voxels in the LOTC ROI. We reasoned that if limb-selectivity is repurposed to word-selectivity as children become better readers, limb-selectivity should be negatively linked to reading ability. Results of this analysis revealed a negative correlation between limb-selectivity and reading skills, which was marginally significant after correcting for age. Although no firm conclusions can be drawn from these findings, they raise the possibility that cortical recycling may also occur in the lateral stream. If limb-selectivity in the lateral stream is indeed repurposed to word-selectivity, an open question concerns which limb- and word-selective patches might be involved in the process in this stream. Future longitudinal studies could test if the lateral word-selective patch located on the inferior occipital sulcus (Stigliani et al., 2015; Zhan et al., 2023) may be engaged in cortical recycling.

A related, interesting finding is that selectivity to pseudowords increased from childhood to adulthood in the left LOTC, but that no such increase was observed for real words. In fact, while adults showed higher activity to pseudowords compared to real words, children showed similar activity across different types of word stimuli. In adults, higher activity for pseudowords compared to real words (Dehaene et al., 2002) and for low-frequency words compared to high-frequency words (Kronbichler et al., 2004; Woolnough et al., 2021) in word-selective regions has been reported repeatedly. It is thought that this pattern of results may reflect easier activation of representations of frequently compared to infrequently used words (Kronbichler et al., 2004) and that pseudowords evoke the strongest activity because they require the longest search times in the neural lexicon (and ultimately do not match any representations in there; Woolnough et al., 2021). Thus, these results raise the possibility that activity for pseudowords in children is lower compared to adults because their neural lexicon is smaller and may lead to shorter search times.

Our analyses also shed light on the specificity of the decrease of limb-selectivity: We examined how the development of limb-selectivity relates to (i) other body stimuli and (ii) tools, which are manipulated by hands. While a prior developmental study suggested a decrease in size of the lateral right EBA from childhood to adulthood (Peelen et al., 2009), our results reveal no evidence for a developmental change for whole-body stimuli, mirroring previous work on the ventral stream (Nordt et al., 2021; Peelen et al., 2009). Interestingly, we found that selectivity to pens, one of the tool categories, decreased from childhood to adulthood both in the ventral and in the lateral ROI. While this result should be interpreted with caution, as the number of tool-selective voxels was overall small and no development was observed for shovels, it may reflect differences in viewing frequency of this category across groups: Pens are frequently used and seen in every-day tasks at school and possibly less so in adulthood. In addition, a longitudinal study tracking development of high-level visual regions in children over the first year of school showed that word-selective voxels emerged in cortex with a prior selectivity to tools (Dehaene-Lambertz et al., 2018). In sum, these findings suggest the possibility that limb- and tool-selective regions may share some developmental processes, which is in line with the finding that tool- and hand-selective regions partially overlap (Bracci et al., 2012; Pillet et al., 2024).

Our results also raise the question whether the decrease in limb-selectivity comes with behavioural consequences. For example, previous studies showed that increased face-selectivity was directly linked to higher performance on a face recognition task (Golarai et al., 2007) and word-selectivity in VTC to reading skills (Kubota et al., 2019). Yet, testing what may be the behavioural consequences of the decrease of limb-selectivity is complex. One reason for this is that the information derived from perceiving body parts like limbs plays a crucial role in several social-cognitive processes (Downing & Peelen, 2016), which include gathering information on someone’s emotional state (Blythe et al., 2023), emphasizing speech, or to identify (object-directed) actions (Wurm & Schubotz, 2017) or goals (Woodward, 1998). Thus, future research is required to explore whether and how changes in limb-selectivity may relate to these different kinds of behaviours.

The present results also enhance our understanding of the development of different visual streams more broadly. Our results indicate that limb-selective regions develop similarly across the ventral and the lateral stream. As such, they differ from previous research showing a differential development of category-selective regions across streams (Golarai et al., 2007). Specifically, this research has shown that ventral, but not lateral face-selective regions expand with age, and the size of the ventral face-selective region was linked to face recognition skills (Golarai et al., 2007). In combination, these results suggest that for some category representations, development is stream-specific, while this is not the case for others. Nonetheless, it is possible that the developmental trajectories of limb- and body-selective regions in the lateral and ventral streams may differ in other developmental periods.

The age range of the present sample of children (10–12 years) is relevant when interpreting the timing of the observed developmental effects. First, these findings are consistent with evidence for a prolonged maturation of high-level visual cortex that extends beyond the age of 10 (Golarai et al., 2010; Nordt et al., 2021, 2023), and with research showing that neural systems supporting social attention and interactive processing continue to refine across middle childhood and adolescence (Oberwelland et al., 2016; Redcay & Warnell, 2018).Second, these results underscore the importance of studying younger developmental samples. For example, fMRI work in infants suggests that the ventral body-selective area (FBA) may emerge later than the lateral body-selective area (EBA; Kosakowski et al., 2022). Future longitudinal research spanning infancy and the preschool years, and using both static and dynamic stimuli (Pitcher et al., 2019), will be crucial for delineating the developmental trajectories of limb- and body-selective regions across visual streams in early development.

## Data statement

Aggregated data and code to generate the main figures will be made publicly available on GitHub. Raw (f)MRI data cannot be made publicly available because sharing is restricted under Institutional Review Board (IRB) regulations.

## Acknowledgements

We thank Eva Broszy, Theresa Heinen, Lilly Baksa and members of the Imaging Core Facility (ICF) at Forschungszentrum Jülich for help with data collection.

## Author Contributions

S.C. conducted and coordinated the project, collected and analysed the data and wrote the manuscript. L.K. collected part of the data and corrected and reviewed the manuscript. S.D.Y. corrected and reviewed the manuscript. M.N. was responsible for funding and designing of the project, contributed to the analysis pipeline and corrected and reviewed the manuscript.

