## SupplementaryFiguresAndTables for "Limb-Selective Regions in the Lateral Temporal Lobe Shrink from Childhood to Adulthood"

**Supplementary Figures**

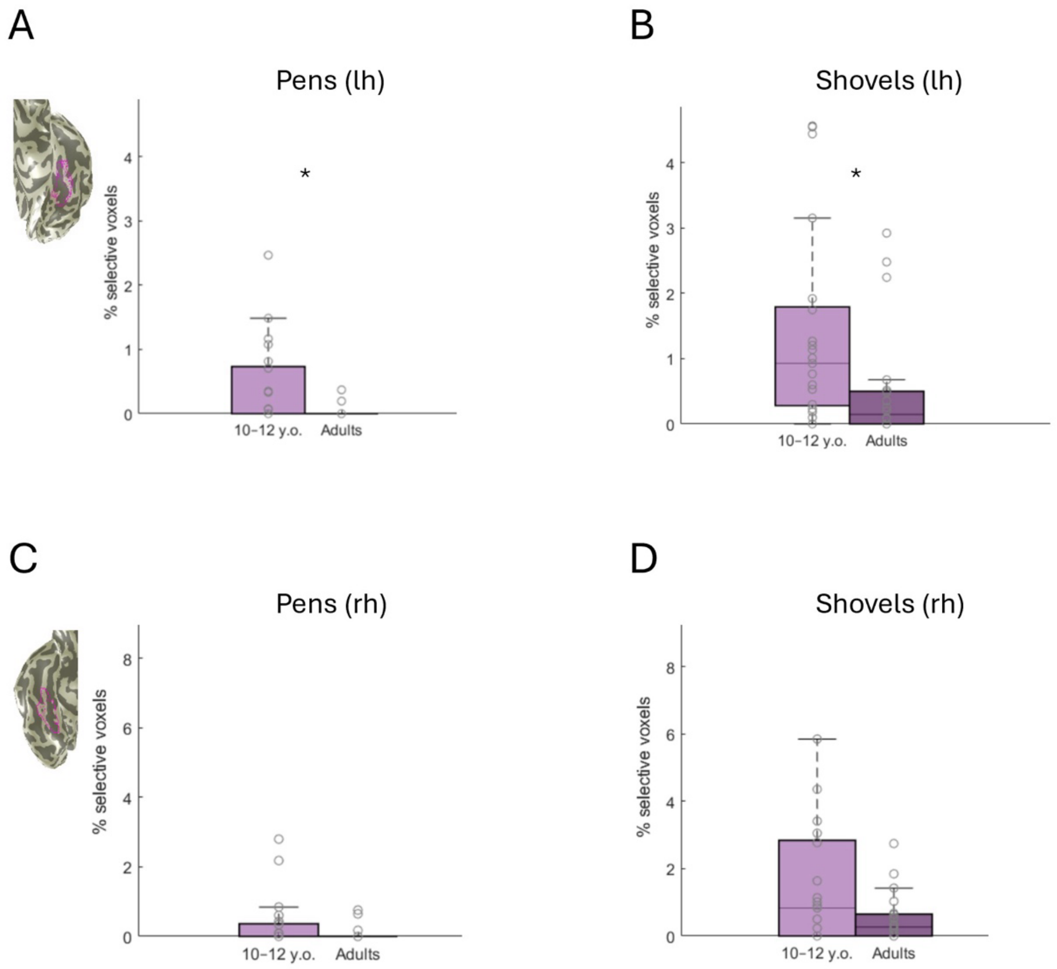


**Supplementary Figure 1. A.** Number of voxels in % out of all voxels in the FG3 ROI in the left hemisphere reacting selectively to pens. Inset: Example of a ventral FG3 ROI in the left hemisphere of an 11-year-old participant. The horizontal line in each box denotes the median value. Whiskers extend to the most extreme data points that do not qualify as outliers. Asterisks indicate a significant group difference ($p<0.05$). **B.** Same as A, but for shovels. **C.** Same as A, but for the right hemisphere. Inset: Example of a ventral FG3 ROI in the right hemisphere of an 11-year-old participant. **D.** Same as A, but for the right hemisphere and for shovels.

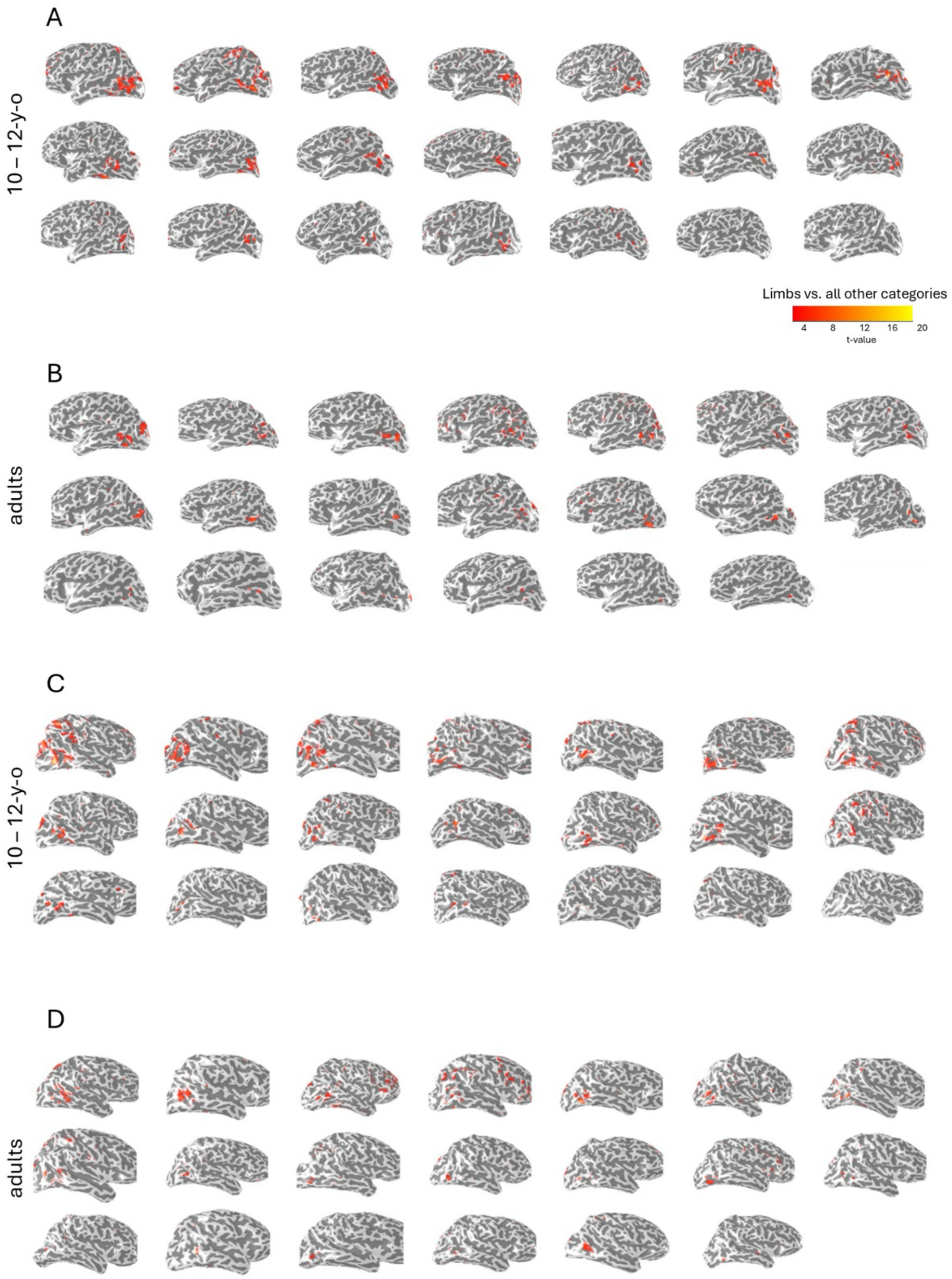


**Supplementary Figure 2. A.** Left inflated cortical surfaces of children sorted by the number of voxels reacting selectively to limbs relative to all voxels in the left LOTC ROI. Limb-selective patches are shown in red. **B.** Same as A, but for adults. **C.** Same as A, but for the right hemisphere. **D.** Same as A, but for the right hemisphere in adults.

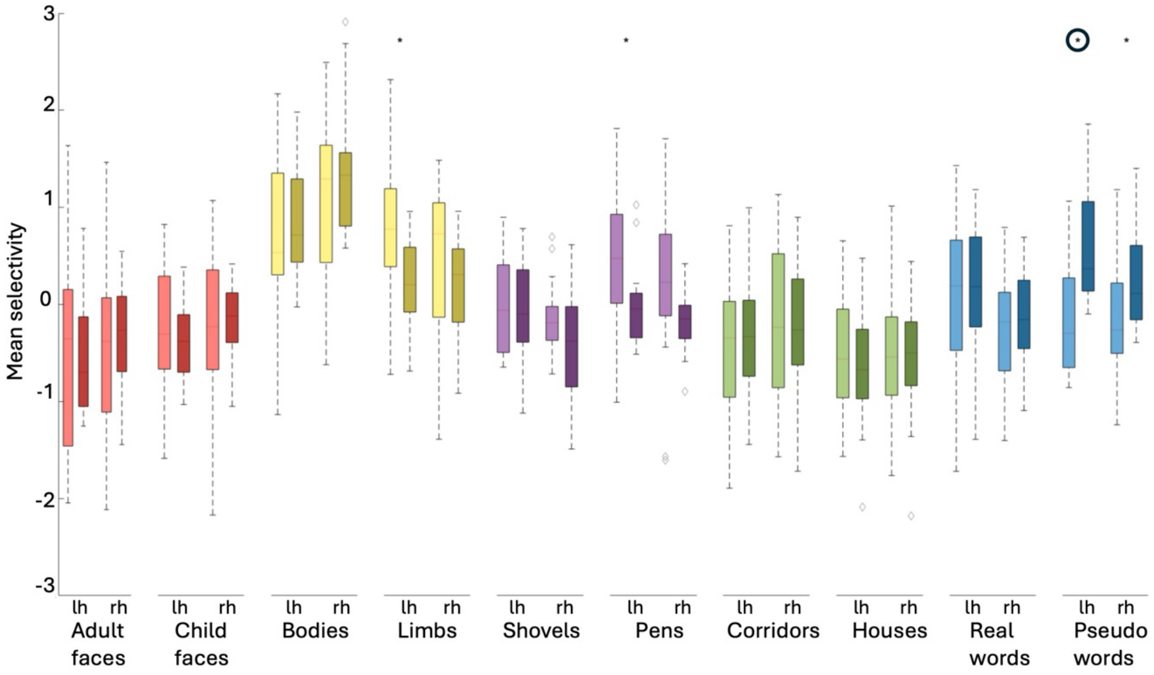


**Supplementary Figure 3.** Mean Selectivity in t-values across all voxels in the LOTC ROI. For each category, selectivity was computed by contrasting the responses to the given category to those of all other categories in the experiment. The horizontal line in each box denotes the median value. Whiskers extend to the most extreme data points that do not qualify as outliers. Diamonds are outliers, defined as lying beyond 1.5 interquartile ranges from the first or third quartile. Asterisks indicate a significant group difference ($p<0.05$). Circles around the asterisks indicate that the effect survives correction for multiple comparisons across the 10 categories.

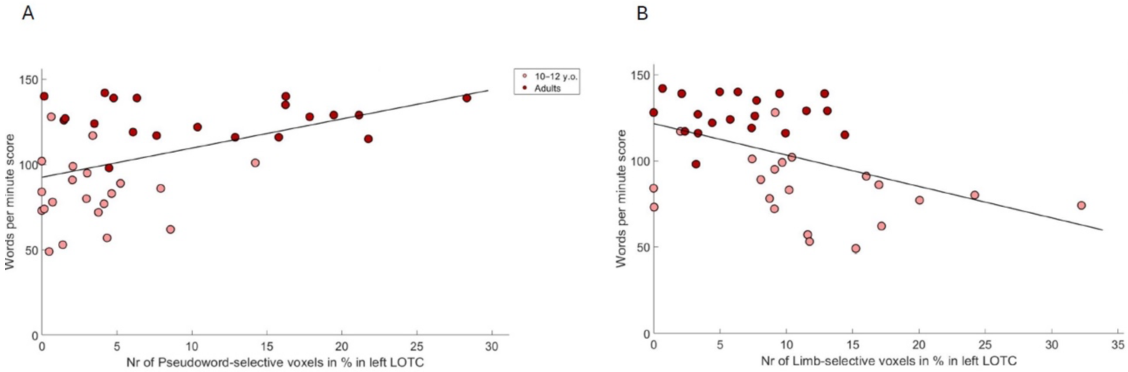


**Supplementary Figure 4. A.** Correlation between reading ability (the number of words read within one minute) and the relative number of voxels reacting selectively to pseudowords in the left LOTC ROI (r = 0.454; p = 0.0029). The correlation remains significant when age is accounted for (controlled for age r = 0.349; p = 0.0275). **B.** Correlation between reading ability (the number of words read within one minute) and the relative number of voxels reacting selectively to limbs in the left LOTC ROI (r = -0.499; p = 0.0032). When age is controlled for, the correlation is trending (controlled for age r = 0.287; p = 0.0723).

### Supplementary Tables

**Supplementary Table 1.** Levene Test in the left (lh) and right (rh) hemispheres in the FG4 ROIs for the relative number of voxels selective to the respective category. Asterisks indicate that the Levene Test yielded significant results. If this was the case, Welch's T-Test was performed for the respective categories.

| Category | Hemisphere | Levene Test (p-Value) |
|---|---|---|
| Adult Faces | lh | 0.023* |
| Adult Faces | rh | 0.116 |
| Child Faces | lh | 0.04 |
| Child Faces | rh | 0.558 |
| Bodies | lh | 0.043* |
| Bodies | rh | 0.024* |
| Limbs | lh | 0.002* |
| Limbs | rh | 0.000* |
| Shovels | lh | 0.171 |
| Shovels | rh | 0.209 |
| Pens | lh | 0.000* |
| Pens | rh | 0.001* |
| Corridors | lh | 0.014* |
| Corridors | rh | 0.462 |
| Houses | lh | 0.09 |
| Houses | rh | 0.12 |
| Words | lh | 0.923 |
| Words | rh | 0.351 |
| Pseudowords | lh | 0.02* |
| Pseudowords | rh | 0.004* |

**Supplementary Table 2.** Independent samples t-tests or Welch's t-test determining whether the number of category-selective voxels (in % out of all voxels in the ROI) differ between children and adults in the FG4 ROI in the left hemisphere. Df=degrees of freedom. Bold p-values indicate the use of Welch's t-test. Asterisks indicate significant results of the t-tests.

| Category | Hemisphere | df | T | p | FDR corrected p |
|---|---|---|---|---|---|
| Adult Faces | lh | 39 | -0.692 | **0.493** | 0.596 |
| Child Faces | lh | 39 | -0.7 | **0.488** | 0.596 |
| Bodies | lh | 39 | 0.748 | **0.459** | 0.596 |
| Limbs | lh | 39 | 2.631 | **0.012*** | 0.05 |
| Shovels | lh | 39 | -0.673 | 0.505 | 0.561 |
| Pens | lh | 39 | 2.876 | **0.007*** | 0.051 |
| Corridors | lh | 39 | 1.507 | **0.14** | 0.399 |
| Houses | lh | 39 | 0.851 | 0.4 | 0.561 |
| Words | lh | 39 | -0.185 | 0.855 | 0.854 |
| Pseudowords | lh | 39 | -2.542 | **0.015*** | 0.07 |

**Supplementary Table 3.** Same as Supplementary Table 2 but for the right hemisphere.

| Category | Hemisphere | df | T | p | FDR corrected p |
|---|---|---|---|---|---|
| Adult Faces | rh | 39 | -1.205 | 0.235 | 0.416 |
| Child Faces | rh | 39 | -0.882 | 0.383 | 0.479 |
| Bodies | rh | 39 | 0.671 | **0.506** | 0.596 |
| Limbs | rh | 39 | 2.092 | **0.043*** | 0.215 |
| Shovels | rh | 39 | -0.33 | 0.744 | 0.743 |
| Pens | rh | 39 | 1.975 | **0.055** | 0.222 |
| Corridors | rh | 39 | -0.477 | 0.636 | 0.707 |
| Houses | rh | 39 | -1.168 | 0.25 | 0.416 |
| Words | rh | 39 | -0.938 | 0.354 | 0.479 |
| Pseudowords | rh | 39 | -1.831 | **0.075** | 0.249 |

**Supplementary Table 4.** Levene Test in the left (lh) and right (rh) hemispheres in the LOTC ROIs for the relative number of voxels selective to the respective category. Asterisks indicate that the Levene Test yielded significant results. If this was the case, Welch's T-Test was performed for the respective categories.

| Category | Hemisphere | Levene Test (p-Value) |
|---|---|---|
| Adult Faces | lh | 0.553 |
| Adult Faces | rh | 0.808 |
| Child Faces | lh | 0.549 |
| Child Faces | rh | 0.139 |
| Bodies | lh | 0.126 |
| Bodies | rh | 0.115 |
| Limbs | lh | 0.102 |
| Limbs | rh | 0.136 |
| Shovels | lh | 0.796 |
| Shovels | rh | 0.849 |
| Pens | lh | 0.000* |
| Pens | rh | 0.009* |
| Corridors | lh | 0.498 |
| Corridors | rh | 0.525 |
| Houses | lh | 0.353 |
| Houses | rh | 0.727 |
| Words | lh | 0.553 |
| Words | rh | 0.29 |
| Pseudowords | lh | 0.000* |
| Pseudowords | rh | 0.308 |

**Supplementary Table 5.** Independent samples t-tests or Welch's t-tests determining whether the number of category-selective voxels (in % out of all voxels in the ROI) differ between children and adults in the LOTC ROI in the left hemisphere. Df=degrees of freedom. Bold p-values indicate the use of Welch's t-test. Asterisks indicate significant results of the t-tests.

| Category | Hemisphere | df | T | p | FDR corrected p |
|---|---|---|---|---|---|
| Adult Faces | lh | 39 | 0.191 | 0.85 | 0.875 |
| Child Faces | lh | 39 | 0.165 | 0.87 | 0.875 |
| Bodies | lh | 39 | -0.341 | 0.19 | 0.475 |
| Limbs | lh | 39 | 2.739 | 0.009* | 0.044 |
| Shovels | lh | 39 | 0.159 | 0.875 | 0.875 |
| Pens | lh | 39 | 2.599 | **0.013** | 0.088 |
| Corridors | lh | 39 | -0.677 | 0.502 | 0.875 |
| Houses | lh | 39 | -0.394 | 0.696 | 0.875 |
| Words | lh | 39 | -0.454 | 0.652 | 0.875 |
| Pseudowords | lh | 39 | -3.964 | **0.0003** | 0.006 |

**Supplementary Table 6.** Same as Supplementary Table 5, but for the right hemisphere.

| Category | Hemisphere | df | T | p | FDR corrected p |
|---|---|---|---|---|---|
| Adult Faces | rh | 39 | -0.497 | 0.622 | 0.778 |
| Child Faces | rh | 39 | 0.997 | 0.325 | 0.759 |
| Bodies | rh | 39 | -1.052 | 0.96 | 0.957 |
| Limbs | rh | 39 | 2.001 | 0.052 | 0.386 |
| Shovels | rh | 39 | 0.246 | 0.807 | 0.897 |
| Pens | rh | 39 | 1.815 | **0.077** | 0.309 |
| Corridors | rh | 39 | 0.710 | 0.482 | 0.778 |
| Houses | rh | 39 | 0.889 | 0.38 | 0.759 |
| Words | rh | 39 | 0.554 | 0.583 | 0.778 |
| Pseudowords | rh | 39 | -1.418 | 0.164 | 0.547 |

**Supplementary Table 7.** Levene Test in the left (lh) and right (rh) hemispheres in the LOTC ROIs for the absolute number of voxels selective to the respective category. Asterisks indicate that the Levene Test yielded significant results. If this was the case, Welch's T-Test was performed for the respective categories.

| Category | Hemisphere | Levene Test (p-Value) |
|---|---|---|
| Adult Faces | lh | 0.337 |
| Adult Faces | rh | 0.493 |
| Child Faces | lh | 0.434 |
| Child Faces | rh | 0.1 |
| Bodies | lh | 0.041* |
| Bodies | rh | 0.03* |
| Limbs | lh | 0.04* |
| Limbs | rh | 0.067 |
| Shovels | lh | 0.862 |
| Shovels | rh | 0.789 |
| Pens | lh | 0.000* |
| Pens | rh | 0.004* |
| Corridors | lh | 0.551 |
| Corridors | rh | 0.678 |
| Houses | lh | 0.268 |
| Houses | rh | 0.506 |
| Words | lh | 0.261 |
| Words | rh | 0.284 |
| Pseudowords | lh | 0.002* |
| Pseudowords | rh | 0.2 |

**Supplementary Table 8.** Independent samples t-tests or Welch's t-tests determining whether the absolute number of voxels in the ROI differs between children and adults in the LOTC in the left hemisphere. Df=degrees of freedom. Bold p-values indicate the use of Welch's t-test. Asterisks indicate significant results of the t-tests.

| Category | Hemisphere | df | T | p | FDR corrected p |
|---|---|---|---|---|---|
| Adult Faces | lh | 39 | 0.359 | 0.722 | 0.932 |
| Child Faces | lh | 39 | 0.206 | 0.838 | 0.932 |
| Bodies | lh | 39 | -0.086 | **0.932** | 0.932 |
| Limbs | lh | 39 | 2.879 | **0.006*** | 0.056 |
| Shovels | lh | 39 | 0.104 | 0.918 | 0.932 |
| Pens | lh | 39 | 2.778 | **0.008*** | 0.056 |
| Corridors | lh | 39 | -0.539 | 0.593 | 0.932 |
| Houses | lh | 39 | -0.477 | 0.636 | 0.932 |
| Words | lh | 39 | -0.569 | 0.573 | 0.932 |

| | | | | | |
|---|---|---|---|---|---|
| Pseudowords | lh | 39 | -3.663 | **0.001*** | 0.015 |

**Supplementary Table 9.** Same as Supplementary Table 8, but for the right hemisphere.

| **Category** | **Hemisphere** | **df** | **T** | **p** | **FDR corrected p** |
|---|---|---|---|---|---|
| Adult Faces | rh | 39 | -0.172 | 0.864 | 0.932 |
| Child Faces | rh | 39 | 1.055 | 0.298 | 0.745 |
| Bodies | rh | 39 | -0.423 | **0.675** | 0.932 |
| Limbs | rh | 39 | 2.286 | 0.028* | 0.139 |
| Shovels | rh | 39 | 0.281 | 0.78 | 0.932 |
| Pens | rh | 39 | 1.982 | **0.055** | 0.218 |
| Corridors | rh | 39 | 0.913 | 0.367 | 0.815 |
| Houses | rh | 39 | 1.129 | 0.266 | 0.745 |
| Words | rh | 39 | 0.585 | 0.562 | 0.932 |
| Pseudowords | rh | 39 | -1.467 | 0.15 | 0.502 |

**Supplementary Table 10.** Levene Test in the left (lh) and right (rh) hemispheres in the LOTC ROIs for the mean selectivity (in t-values averaged across all voxels in the ROI). Asterisks indicate that the Levene Test yielded significant results. If this was the case, Welch's T-Test was performed for the respective categories.

| **Category** | **Hemisphere** | **Levene Test (p-Value)** |
|---|---|---|
| Adult Faces | lh | 0.04* |
| Adult Faces | rh | 0.026* |
| Child Faces | lh | 0.044* |
| Child Faces | rh | 0.007* |
| Bodies | lh | 0.125 |
| Bodies | rh | 0.066 |
| Limbs | lh | 0.224 |
| Limbs | rh | 0.099 |
| Shovels | lh | 0.618 |
| Shovels | rh | 0.026* |
| Pens | lh | 0.028* |
| Pens | rh | 0.01* |
| Corridors | lh | 0.863 |
| Corridors | rh | 0.3 |
| Houses | lh | 0.703 |
| Houses | rh | 0.51 |
| Words | lh | 0.354 |
| Words | rh | 0.33 |

| | | |
|---|---|---|
| Pseudowords | lh | 0.654 |
| Pseudowords | rh | 0.787 |

**Supplementary Table 11.** Independent samples t-tests or Welch's t-tests determining whether the mean selectivity (in t-values averaged across all voxels in the ROI) differs between children and adults in the LOTC ROI in the left hemisphere. Df=degrees of freedom. Bold p-values indicate the use of Welch's t-test. Asterisks indicate significant results of the t-tests.

| Category | Hemisphere | df | T | p | FDR corrected p |
|---|---|---|---|---|---|
| Adult Faces | lh | 39 | 0.369 | **0.714** | 0.841 |
| Child Faces | lh | 39 | 0.049 | **0.961** | 0.961 |
| Bodies | lh | 39 | -0.743 | 0.462 | 0.841 |
| Limbs | lh | 39 | 2.974 | 0.005* | 0.05 |
| Shovels | lh | 39 | 0.58 | 0.566 | 0.841 |
| Pens | lh | 39 | 2.21 | **0.01*** | 0.167 |
| Corridors | lh | 39 | -0.727 | 0.472 | 0.841 |
| Houses | lh | 39 | 0.641 | 0.525 | 0.841 |
| Words | lh | 39 | -0.371 | 0.713 | 0.841 |
| Pseudowords | lh | 39 | -4.55 | 0.0003* | 0.001 |

**Supplementary Table 12.** Same as Supplementary Table 11, but for the right hemisphere.

| Category | Hemisphere | df | T | p | FDR corrected p |
|---|---|---|---|---|---|
| Adult Faces | rh | 39 | -0.405 | **0.688** | 0.841 |
| Child Faces | rh | 39 | -0.453 | **0.653** | 0.841 |
| Bodies | rh | 39 | -1.046 | 0.302 | 0.755 |
| Limbs | rh | 39 | 1.418 | 0.164 | 0.511 |
| Shovels | rh | 39 | 1.37 | **0.179** | 0.511 |
| Pens | rh | 39 | 1.906 | **0.08** | 0.256 |
| Corridors | rh | 39 | 0.253 | 0.801 | 0.844 |
| Houses | rh | 39 | 0.31 | 0.758 | 0.842 |
| Words | rh | 39 | -0.758 | 0.453 | 0.841 |
| Pseudowords | rh | 39 | -2.372 | 0.023* | 0.151 |
